# Comparative proteome signatures of trace samples by multiplexed Data-Independent Acquisition

**DOI:** 10.1101/2021.02.11.430601

**Authors:** Claudia Ctortecka, Gabriela Krššáková, Karel Stejskal, Josef M. Penninger, Sasha Mendjan, Karl Mechtler, Johannes Stadlmann

**Affiliations:** Research Institute of Molecular Pathology (IMP), Vienna BioCenter (VBC), Campus-Vienna-Biocenter 1, 1030 Vienna, Austria; Institute of Molecular Biotechnology of the Austrian Academy of Sciences (IMBA), Vienna BioCenter (VBC), Dr. Bohr-Gasse 3, 1030 Vienna, Austria; The Gregor Mendel Institute of Molecular Plant Biology of the Austrian Academy of Sciences (GMI), Vienna BioCenter (VBC), Dr. Bohr-Gasse 3, 1030 Vienna, Austria; Department of Medical Genetics, Life Sciences Institute, University of British Columbia, Vancouver Campus, 2350 Health Sciences Mall, Vancouver, BC Canada V6T 1Z3; BOKU – University of Natural Resources and Life Sciences, Muthgasse 18, A 1190 Vienna, Austria

## Abstract

Single cell transcriptomics has revolutionized our understanding of basic biology and disease. Since transcript levels often do not correlate with protein expression, it is crucial to complement transcriptomics approaches with proteome analyses at single cell resolution. Despite continuous technological improvements in sensitivity, mass spectrometry-based single cell proteomics ultimately faces the challenge of reproducibly comparing the protein expression profiles of thousands of individual cells. Here, we combine two hitherto opposing analytical strategies, DIA and Tandem-Mass-Tag (TMT)-multiplexing, to generate highly reproducible, quantitative proteome signatures from ultra-low input samples. While conventional, data-dependent shotgun proteomics (DDA) of ultra-low input samples critically suffers from the accumulation of missing values with increasing sample-cohort size, data-independent acquisition (DIA) strategies do usually not take full advantage of isotope-encoded sample multiplexing. We developed a novel, identification-independent proteomics data-analysis pipeline that allows to quantitatively compare DIA-TMT proteome signatures across hundreds of samples independent of their biological origin, and to identify cell types and single protein knockouts. We validate our approach using integrative data analysis of different human cell lines and standard database searches for knockouts of defined proteins. These data establish a novel and reproducible approach to markedly expand the numbers of proteins one detects from ultra-low input samples, such as single cells.

## Main

Single-cell proteomics aims at assessing protein expression within individual cells with far-reaching opportunities for a better understanding of fundamental biology or disease states. Currently, protein analysis at single cell resolution is still largely antibody based, therefore relying on the availability of such. This not only greatly limits the throughput of these techniques, but also requires pre-formed hypotheses (i.e. flow cytometry and mass cytometry). At present, mass spectrometry-based proteomics is the only viable technology for discovery and hypothesis-free protein analysis.

While the comprehensive proteomic characterization of individual mammalian cells is still limited by the sensitivity of current MS/MS-based workflows, the concept of multiplexed shotgun proteomics analyses of individual cells in conjunction with a highly abundant, congruent carrier proteome has been seminal to the field.^1^ The use of established *in vitro* stable-isotope labeling techniques (e.g. TMT) not only increases precursor- and fragment-ion abundances for peptide identification and quantification from ultra-low input samples, but also increases sample throughput. Currently, such multiplex single-cell proteomics workflows have allowed for the quantitative analysis of up to 13 barcoded single cells in one analytical run.^2^

Nevertheless, paralleling state-of-the-art transcriptomic datasets, single-cell proteomics ultimately faces the challenge to comparatively analyze hundreds or even thousands of ultra-low input proteomics samples.^3–5^ Such sample sizes vastly exceed the capacities of any currently available MS multiplexing technology^6^. Merging large numbers of individual quantitative shotgun proteomics (DDA) files into one dataset, often entails that a considerable number of peptides are not reliably identified in all analytical runs.^7^ This method-intrinsic accumulation of “missing values” greatly limits the use of DDA strategies for the comparative analysis of protein levels in large sample numbers, as are necessary for reproducible single cell proteomics, which is currently addressed by various computational data-imputation or “match-between runs” methods.^8–10^

By contrast, data-independent acquisition (DIA) regimes, which subject all precursor ions within a defined m/z window to MS/MS analysis, have been shown to allow for the robust quantification of protein expression, even across extremely large sample cohorts.^11^ Recently, DIA strategies were further extended to sequentially windowed DIA-schemes (SWATH), specifically designed to cover all theoretical mass-spectra and to thereby provide deep proteome coverage.^12,13^ To develop a scalable high-throughput data-acquisition strategy for comparative single-cell proteomics, we combined *in vitro* multiplexing strategies for MS/MS based quantification (i.e. TMT10plex ™ Isobaric Label Reagent Set) and small window DIA data-acquisition regimes (i.e. m/z = 6 Th) for the analysis of ultra-low protein amounts.

### DIA-TMT provides reproducible, quantitative proteome signatures

We first acquired DDA and DIA MS/MS-data sets in technical triplicates of TMT10plex-labelled tryptic digests derived from two human cell lines, serially diluted to total peptide amounts similar to those expected for single mammalian cells (i.e. 0.3 ng and lower) (illustrated in Fig. 1a). Two commercially available tryptic digests of whole human cell lysates (i.e. HeLa and K562) were labeled with all available TMT10plex-tags, respectively. The labeled standards were then combined into three different two-proteome mixes equipped with distinct sets of isobaric tags (detailed in the methods section). Each of the three mixes were then analyzed at four different peptide concentrations, i.e. 10, 5, 1 and 0.5 ng total peptide, by nano-HPLC-ESI-MS/MS, using an Orbitrap Exploris™ 480 Mass Spectrometer, fitted with FAIMS (for detailed acquisition parameters see methods section).

**Figure 1:**
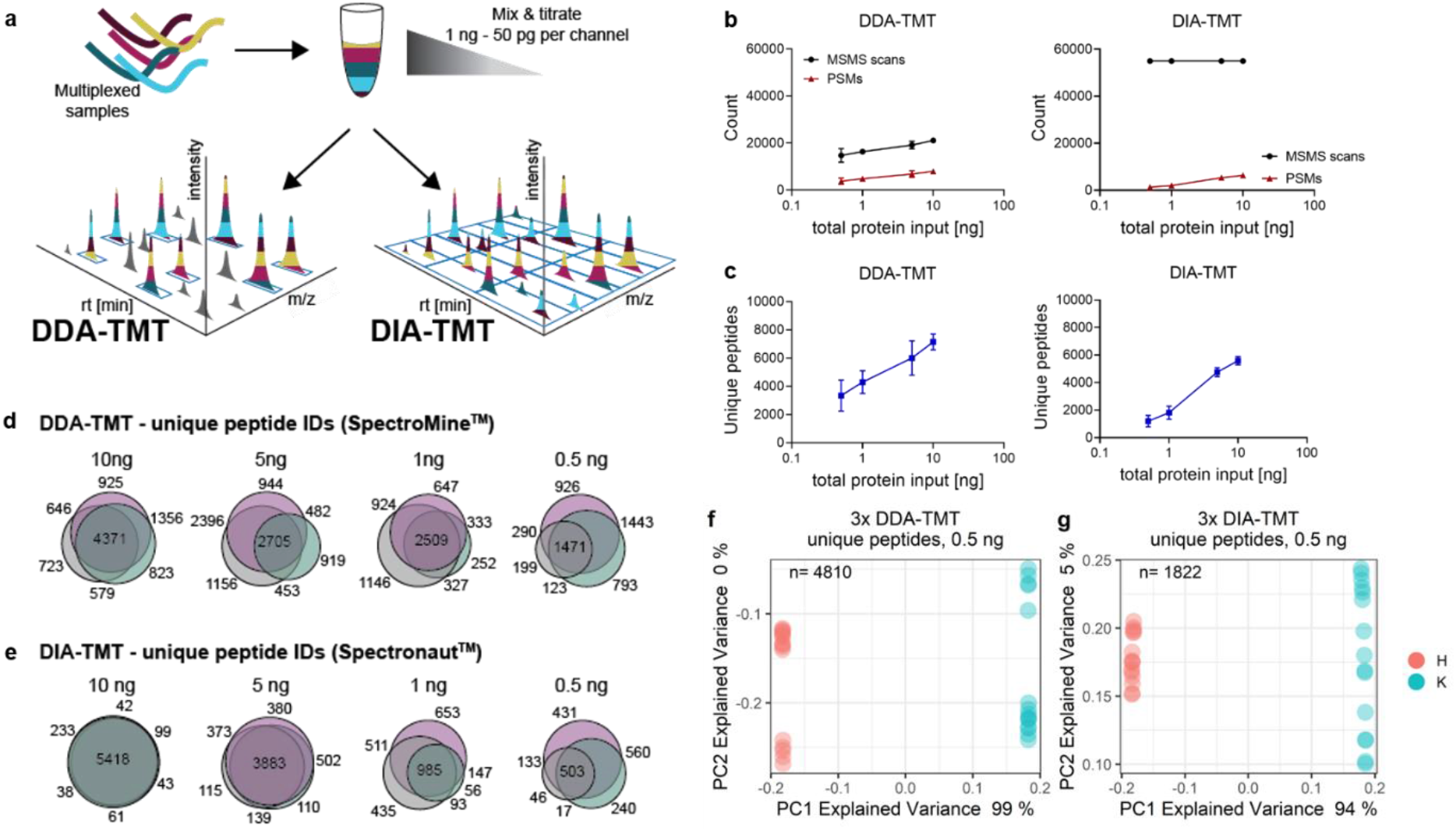
Evaluation of DIA and DDA schemes. **(a)** Graphical abstract of the workflow, **(b)** comparison of DDA and DIA experiments, showing the numbers of MS/MS scans (black), PSMs (brown) and **(c)** unique peptides (blue) using Spectromine™ database search (DDA) or Spectronaut™ library search (DIA) at various peptide input levels (i.e. 0.5 ng, 1 ng, 5 ng, 10 ng). Mean and standard deviations are shown. Venn Diagrams of unique peptide sequences identified across three technical replicates at indicated peptide input of **(d)** DDA and **(e)** DIA data (mix 1 = purple, mix 2 = grey, mix 3 = blue). PCA clustering of three analytical runs at 0.5 ng total peptide input acquired with **(f)** DDA and **(g)** DIA acquisition schemes, aggregated via the standard identification-dependent method. The two different colors represent channel loadings, n= number of unique peptides included in PCA. H = Hela, K = K562.

Manual inspection of the raw DIA MS/MS-data confirmed a robust and input-independent number of MS/MS scans. In DDA, however, the number of MS/MS scans markedly decreased with lower amounts of peptide input (Fig. 1b). Nevertheless, subsequent peptide identification by dedicated MS/MS search engines (i.e. Spectronaut™ for DIA and Spectromine™ for DDA data) yielded twice as many peptide-spectrum matches (PSMs) and unique peptide identifications from DDA data, even at lower concentrations, when compared to corresponding DIA data. At 0.5 ng total peptide input we identified 1.302 and 3.843 PSMs or 1.202 and 3.348 unique peptides in the DIA and DDA acquisition modes, respectively. The 1 ng total peptide input samples yielded 1.982 and 4.856 PSMs or 1.817 and 4.296 unique peptides from DIA and DDA data, respectively (Fig. 1b-c). These findings demonstrate that, despite steadily generating an enormous amount of MS/MS data, especially in DIA, the number of identified peptides markedly decreases with lower sample input.

To evaluate scalability of DIA strategies for peptide identification from multiple samples, we merged quantitation data from several analytical runs, based on peptide identity. For this, we only considered peptides that were consistently identified in all three technical replicates at a given total peptide input level. As expected, at 10 ng total peptide input, we observed that DIA indeed provided more consistent peptide identifications across multiple analytical runs (i.e. > 90 % central overlap), when compared to DDA (i.e. < 50 % central overlap) (Fig. 1d-e). Importantly, this key benefit of DIA strategies was gradually lost with decreasing peptide input (Fig. 1d), presumably because of decreasing total ion current. This data shows that, like DDA, also DIA fails to provide consistent peptide identifications across multiple ultra-low input samples, therefore also giving rise to detrimental amounts of missing values.

Next, we assessed the quantitative information obtained by the two MS/MS-data acquisition strategies. We first employed a conventional, identification-based TMT-data analysis workflow with standard missing value imputation and analyzed unique peptide-derived TMT reporter-ion intensities by principal component analysis (PCA) for the two human cell lines HeLa and K562. Although different numbers of data points (i.e. DDA: 4810, DIA: 1822) were used for PCA, both unique peptide sets allowed a distinction between the two human cell lines via the first two principal components, even at 0.5 ng total protein level. For both datasets the first principal component (PC), which is displayed on the x-axis separates the cell lines with over 90% explained variance, while the second PC representing close to 0% of the variance discriminates between the analytical runs (y-axis) (Fig. 1f-g). This data indicates that through identification-dependent aggregation of DDA and DIA data the biological variance is the main characteristic (Fig. 1f-g).

### Identification-independent analysis of DIA-TMT data precludes “missing data”

We hypothesized that the generation of comprehensive, quantitative proteome signatures rather than peptide-based profiles would allow us to detect subtle expression changes in trace samples. To test this, all datasets were first retention-time (RT) aligned, based on the elution time-points of eight doubly charged peptide precursor-ions, evenly distributed across the entire analytical gradient and consistently detected in all samples (detailed in methods). The RT-aligned DIA data were then aggregated using a dual indexing approach, based on the central m/z of the respective isolation window and the acquisition cycle number as indices. For direct comparison of identification-independent clustering, DDA MS/MS data were binned according to the central m/z of the isolation window (bin-size = 1 Thomson (Th).) and RT (bin-size = 0.2 min; ∼ two times the average chromatographic peak-width), and afterwards matched between files (with a tolerance of +/-1 bin, respectively). The extracted raw TMT-reporter-ion intensities from all aggregated MS/MS spectra, irrespective of peptide identification (using the in-house developed PD-Node IMP-Hyperplex), were then analyzed by PCA. In contrast to the standard identification dependent data-analysis, this identification-independent approach consistently yielded more than 24.000 data-points (Fig. 2a-b) from each DIA run and resulted in PCA clustering of the expected cell populations, for all samples, even at 0.5 ng total protein input (Fig. 2c this should be c, following the flow). Of note, despite less data-points, PCA of the corresponding DDA datasets provided similar clustering of HeLa and K562 cells (Fig. 2d). Thus, although using ultra-low sample amounts, our approach allowed us to correctly identify the cellular source of the protein input.

**Figure 2:**
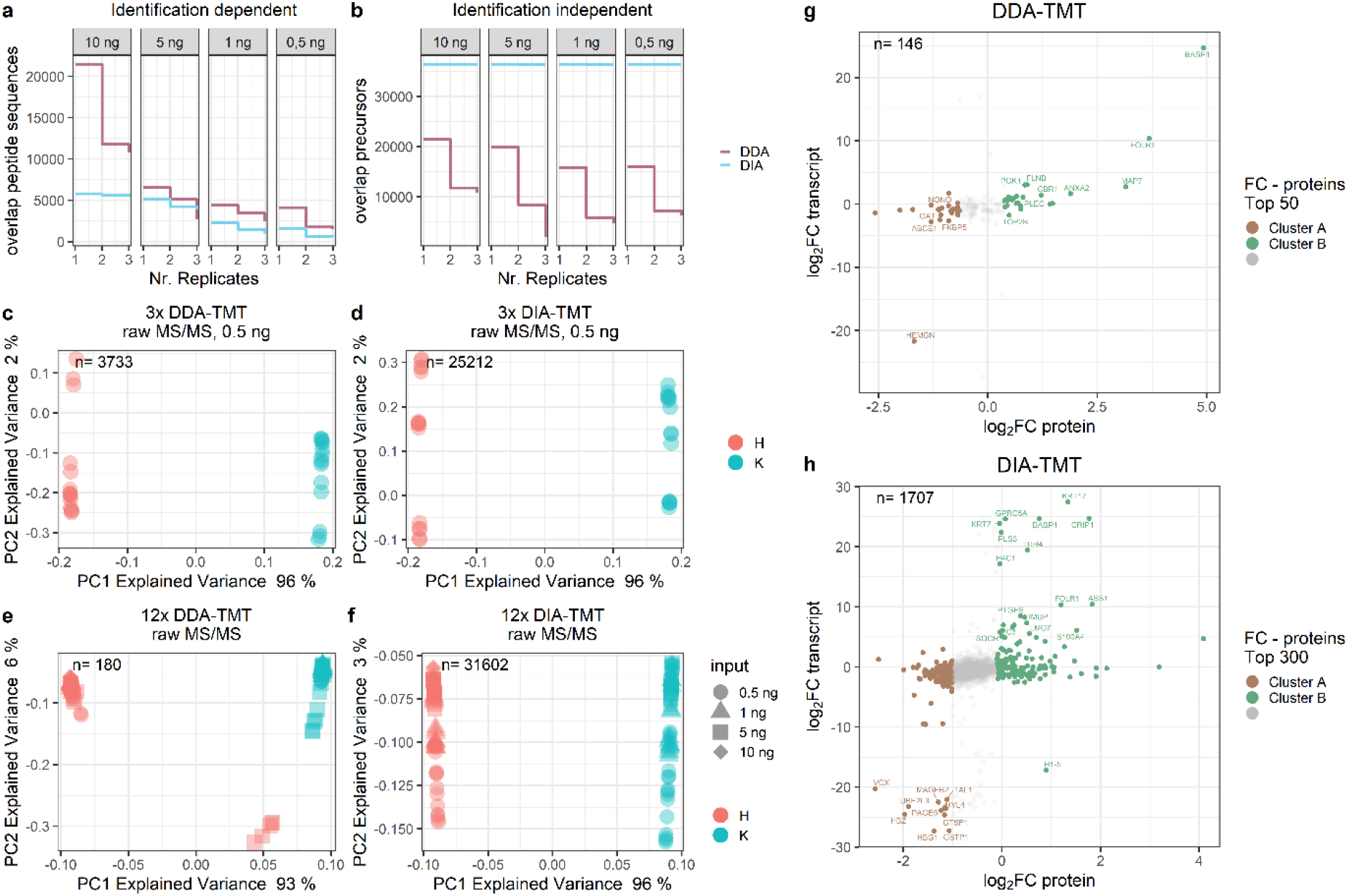
Data aggregation in DDA and DIA acquisition methods. **(a)** Identification-dependent accumulation of non-overlapping peptides and **(b)** identification-independent aggregation showing quantitative data points across 12 analytical runs at decreasing peptide input (i.e. 0.5, 1, 5, 10 ng) using the DDA (purple) and DIA (blue) acquisition schemes. PCA of three analytical runs (30 multiplexed samples) at 0.5 ng total peptide input were acquired with **(c)** DDA or **(d)** DIA identification-independent data aggregation. PCA of twelve analytical runs (120 multiplexed samples) at four peptide inputs (0.5, 1, 5, 10 ng) using **(e)** DDA or **(f)** DIA acquisition schemes, identification-independent aggregated. Samples are colored according to channel loadings and the respective peptide inputs are indicated with different symbols. H = Hela cells, K = K562 cells. n= number of MSMS scans included in PCA. Intersection of transcriptome (FPKM) with proteome data from both **(g)** DDA and **(h)** DIA. Top 300 or 50 proteins are colored according to cluster contributions. Top 60 or 40 proteins are labeled for DIA and DDA, respectively.

We next assessed the scalability of our identification-independent DIA approach. To evaluate whether PCA clustering was independent of the sample amount, we merged all twelve DIA and DDA datasets, respectively. Intriguingly, for both datasets, we observed mix-independent cell type clustering, with over 90% explained variance in PC1 for both DIA and DDA (Fig. 2e-f). However, within the twelve aggregated samples using DDA, we observed a batch effect for one of the samples (i.e. 5 ng, Fig. 2e). Additionally, while precursor-ion sampling stochasticity in DDA markedly reduced the number of MS/MS scans consistently measured and matched across 120 samples to merely 180 scans (Fig. 2e), DIA afforded the consistent accumulation of 31.602 data-points across all samples measured (Fig. 2f). This data highlights that, despite a similar level in the accumulation of missing values when using conventional peptide identification workflows, DIA does indeed generate more robust and highly congruent proteome signatures from larger sample sizes, even at ultra-low input. Most importantly, in contrast to identification dependent data aggregation (Fig. 1g), identification-independent clustering of DIA datasets were found to be largely independent of sample batches (Fig. 2f).

To further validate our workflow to faithfully identify the two cell types, we merged the respective raw MS/MS datasets, re-analyzed them using dedicated MS/MS search engines (Spectromine™ for DDA or Spectronaut™ for DIA), projected the resulting peptide identifications onto the data-tables and calculated fold changes in protein abundance between clusters A and B. This data-analysis allowed us to compare the protein expression levels of 380 and 1.741 proteins by DDA and DIA, respectively. To align differential protein expression to transcriptome data, we then calculated fold changes of fragments per kilobase of transcript per million mapped reads (FPKM) values of HeLa and K562 cells, publicly available via the ENCODE project (GEO accession: GSE33480).^14^ Using the Uniprot database, Ensembl GeneIDs from the transcript data and Protein Accession numbers from Spectromine™ or Spectronaut™, were mapped the expressed genes to our proteomics analysis (DDA: 163 proteins/transcripts; DIA: 1.707 proteins/transcripts). Transcript levels of HeLa and K562 were plotted against protein expression; the top 300 (DIA) or top 50 (DDA) protein fold changes of cluster A and B are shown in Fig. 2g-h. We observed reduced expression ratios in both DDA and DIA protein fold changes as compared to the transcriptomics data (DDA: protein: -3.4 to 3.9, transcript: -27.3 to 24.6; DIA: protein -2.5 to 4.1, transcript: -27.3 to 27.4 in log_2_ space), as expected.^15^ Importantly, fold changes of the transcriptomics results paralleled our proteomics data, confirming that our identification-independent data analysis approach can indeed identify cell type specific clusters. This suggests that our proteome signatures allow for robust clustering and discrimination of cell types, while database re-analysis of these clusters reveals their cellular identity.

### Identification of single-protein knockouts by identification-independent data aggregation

Next, we further investigated whether the DIA acquisition strategy in conjunction with identification-independent data aggregation would also allow discriminating between highly similar single protein knockout cell-lines. Therefore, we generated DIA data sets using the yeast TKO11 standard at four input levels, i.e. 10, 5, 1 and 0.5 ng total peptide in triplicates. This commercially available TMT11-plex labeled tryptic TKO11 yeast standard, comprised of three different single knockout (*met6, his4, ura2*) and wild-type yeast strains, is regularly used for TMT benchmarking experiments.^16^

Intriguingly, after identification independently aggregating three analytical runs at 0.5 ng total peptide input, we observed cell type dependent and batch independent clustering based on almost 16.000 MSMS scans (Fig. 3a). Further, we combined twelve DIA TKO11 runs at four different peptide inputs (0.5, 1, 5, 10 ng) identification independently and observed similar clustering (Fig. 3b). While the combination of multiple analytical runs at the same input capitalizes on the biological differences, the identification independent aggregation impedes clear separation. More specifically, cluster 1 and 2 only comprises the WT strain and the *met6* knockout, respectively, however, cluster 3 combines *his4* and *ura2* knockouts with a trend towards cell type dependent separation (Fig. 3b).

**Figure 3:**
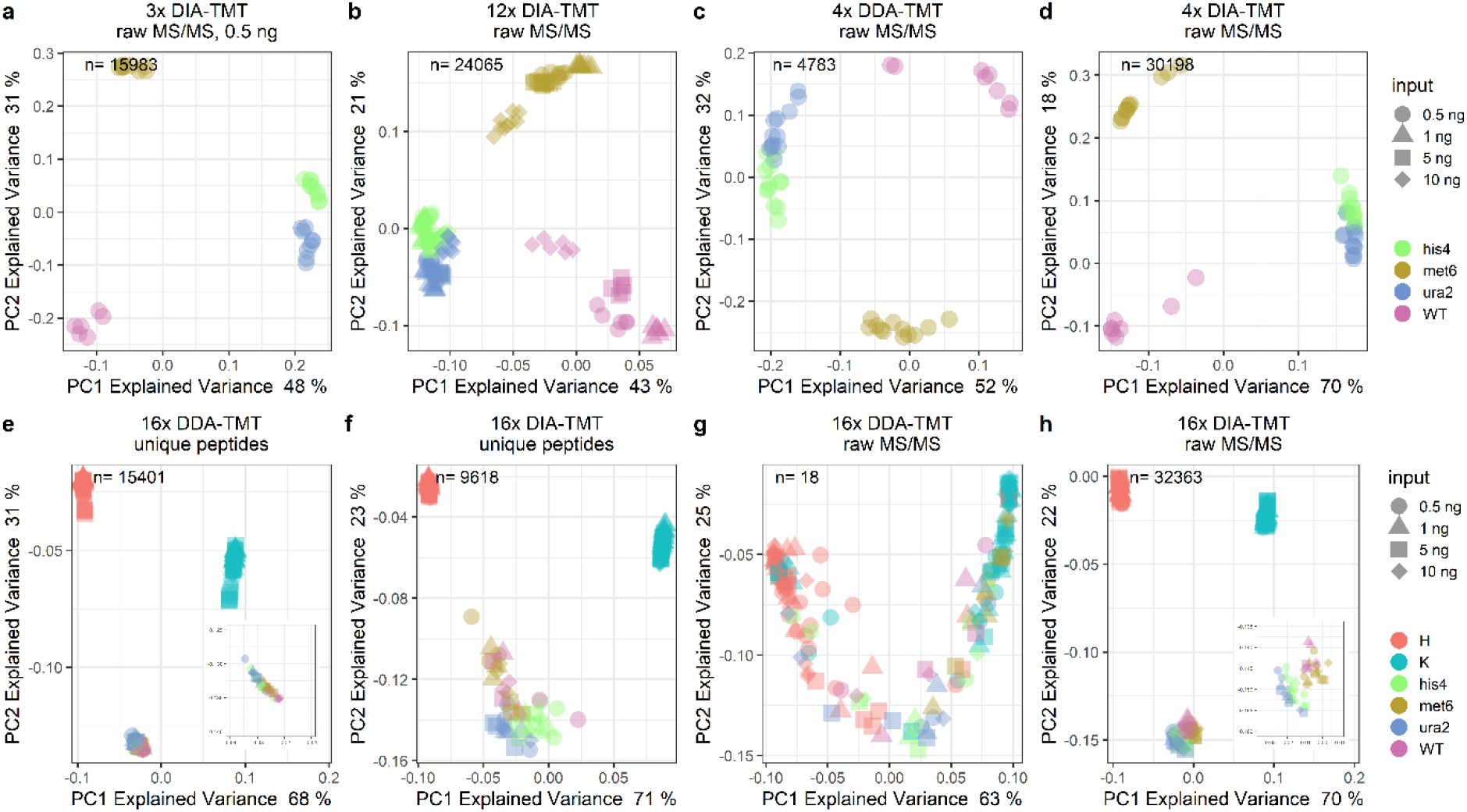
Identification-independent data aggregation allows for the analysis of closely related cell types. PCA of DIA acquisition scheme aggregated using identification independent strategy of yeast TKO11 **(a)** three analytical runs at 0.5 ng peptide input and **(b)** twelve analytical runs at 0.5, 1, 5, or 10 ng peptide. PCA of four analytical yeast TKO11 runs aggregated with identification-independent method at 0.5, 1, 5, or 10 ng peptide inputs. **(c)** DDA and **(d)** DIA. Colors indicate the respective strains (i.e. WT, knockouts: *met6, his4, ura2*). PCA of 16 analytical runs with 0.5, 1, 5, or 10 ng peptide inputs. Standard identification dependent data aggregation with missing value imputation of **(e)** DDA (plus TKO11 zoom) or **(f)** DIA acquisition schemes. n= number of unique peptides included in PCA. PCA of identification-independent aggregation based on 16 analytical runs without imputation of **(g)** DDA or **(h)** DIA data (plus TKO11 zoom). Samples are colored according to channel loadings and the respective peptide input is indicated with different symbols. H = Hela cells, K = K562 cells and the respective TKO11 strains (i.e. WT, knockouts: *met6, his4, ura2*). n= number of MSMS scans included in PCA.

Eventually, to determine the drivers of the observed cell-line clustering we subjected the DIA-TMT TKO11 runs to a standard database search. We identified met6 and ura2 proteins down to 1 ng total peptide input. We then intersected the identified MSMS scans with the loadings of our proteome signature clustering and confirmed these scans as drivers of separation. Even though we did not identify any of the ablated proteins in the 0.5 ng DIA-TMT data, our identification independent data aggregation strategy still allowed for successful cell type dependent clustering. This suggests that standard database searches can be used to infer hypothesis free cell type identifications to the identification-independent proteome signatures.

### DIA-TMT discriminates samples independent of origin or prevalence

We and others^7^ frequently observe file-specific batch effects in DDA (Fig. 2e). While these effects most likely result from stochastic precursor sampling in the course of DDA MS/MS data acquisition, numerous approaches aim at correcting such batch effects pre- or post-acquisition.^7,17,18^ Most importantly, however, using such statistical correction methods, peptides that were only identified in a subset of all analytical runs are extremely prone to over-normalization or exclusion. The need for such data-correction procedures thus critically limits the detection of underrepresented or unexpected cell types (e.g. infiltrated tumor samples) by DDA.

To directly examine the impact of such underrepresented samples we generated data sets analogous to our HeLa and K562 data, yet also including the yeast TKO11 standard at four input levels, i.e. 10, 5, 1 and 0.5 ng total peptide, and post-acquisitively merged DDA and DIA data for analysis via identification-dependent and identification-independent strategies, respectively.

We first evaluated the TKO11 datasets and confirmed cell type-dependent and batch- or input-independent clustering for both DDA and DIA using the identification independent method. Importantly, even upon combination of drastically diverging sample input knockouts *his4* and *ura2* cluster together (Fig. 3a). While identification independent analysis of DDA datasets reduced the overlap to merely 5.000 MSMS scans, DIA analysis retained over 30.000 scans without the need for imputation (Fig. 3c-d). Interestingly, we observed a similar cell-type dependent separation using DDA data when aggregated identification-dependently and based on over 6.000 unique peptides (Supplemental Fig. 2a). Identification dependent analysis of TKO11 DIA analysis, however, drastically reduced the dataset to only 4.246 unique peptides and yielded no appreciable separation based on cellular identity (Supplemental Fig. 2b). Remarkably, based on subsequent database re-analysis we could again infer peptide identifications to the DIA-TMT proteome signatures, which allowed identifying cluster-driving MSMS scans. Expectedly, the strongest driver-scans in both acquisition strategies were post-analytically identified as *met6* knockout peptides (Supplementary Fig. 1a-b). Based on these observations we conclude that the identification independent analysis of DIA and DDA data allows for cell type dependent separation even of single protein knockouts (Fig. 3c-d).

By contrast, using identification dependent data analysis strategies, only up to 17 peptides in both DDA and DIA, were consistently identified in all analytical runs combined. This small group of peptides did not exhibit sufficiently large differences in abundance between the cell types, and thus did not allow for distinct clustering (Supplementary Fig 3a-b). Next, we performed “missing data” imputation based on commonly used Perseus parameters.^19^ Missing values in both, DIA and DDA data, were replaced with random numbers from a normal distribution shifted into the noise. Based on the largely computationally generated quantitative data yeast and human species could be separated via the first PC (Fig. 3e; DDA with 68% and Fig. 3b; DIA with 71% explained variance). Importantly, separation of the individual single protein knockouts was exclusively observed using the DIA acquisition strategy (Fig. 3f).

Furthermore, to detect actual quantitative differences between the samples, we then evaluated our identification-independent data aggregation method on the 16 yeast-human species analytical runs. Strikingly, our DIA data analysis retained critical cell type specific characteristics based on 32.363 quantitative MS/MS scans across 164 samples without the need for any imputation whatsoever (Fig. 3h). Thus, PC1 with 70% explained variance separates the main three species and PC2 with 22% explained variance further differentiates the two species. For direct comparison we also performed PCA analysis of the remaining 18 quantitative data points after identification-independent data aggregation of DDA data and observed no separation, as expected (Fig. 3g).Of note, despite the large variance between two species and the two human cell lines, a zoom into the TKO11 cluster showed that we readily separate between the single yeast mutant strains (Fig. 3h zoom). Thus, our DIA acquisition scheme results in a homogenous dataset can detect small differences within samples.

## Conclusions

Taken together, we here demonstrate that identification independent DIA is scalable and yields meaningful clusters of both, closely related cell types (HeLa vs K562), different composites of distinct species (human versus yeast) and single protein knockout cell lines (TKO11 yeast). Based on standard identification dependent data aggregation methods DIA-TMT performs comparable to DDA-TMT acquisition strategies for ultra-low input samples. However, aggregation of several hundred ultra-low input samples without the accumulation of “missing data” is a major advantage of DIA over standard DDA identification-dependent approaches. Our combination of isobaric multiplexing in DIA acquisition mode allows to generate comprehensive proteome signatures independent of sample origin or input level. We demonstrate that DIA in conjunction with the identification independent data aggregation strategy retains quantitative data across multiple TMT-batches without the need to impute computationally generated values (Fig. 2d, f, Fig. 3 b, d, h). This is in stark contrast to the standard identification dependent method, which drastically reduces quantitative information even in combination with data imputation for both DDA and DIA in the analysis of ultra-low input samples. Furthermore, underrepresented cell populations and their identity can be identified post-acquisition using our method, which is increasingly important when analyzing limited and complex biological samples other than homogenous cell lines. The prospect of species-and cell-type independent analysis would allow for studying diverse samples with no *a priori* knowledge about the specimen. Our novel DIA identification-independent analysis of large numbers and low concentration input samples might thus contribute to a universally applicable workflow for the study of protein expression at single cell resolution.

## Methods

### Sample preparation

Tryptic digests were obtained from Promega (K562, V6951) and Thermo Fisher (HeLa, 88328) were labeled according to manufacturer’s instructions. Briefly, samples were labeled in 100 mM TEAB and 10% ACN for 1 hr at room temperature. Unreacted TMT reagent was quenched with 5% hydroxylamine/HCl for 20 minutes at RT and subsequently mixed corresponding to each TMT10 plex. Mixes were compiled as follows (Mix 1: 126, 127N, 127C, 128N, 128C – K562; 129N, 129C, 130N, 130C, 131 – HeLa; Mix 2: inverted Mix 1; Mix 3: 126, 127C, 128C, 129C, 130C – K562, 127N, 128N, 129N, 130N, 131 – HeLa to exclude any label specific effects.

### LC MS/MS analysis

Samples were measured on a Q Exactive HF-X (Thermo Fisher Scientific) and an Orbitrap Exploris 480 Mass Spectrometer (Thermo Fisher Scientific) with a reversed phase Dionex UltiMate 3000 high-performance liquid chromatography (HPLC) RSLCnano System coupled via a Nanospray Flex ion source equipped with FAIMS Pro. Chromatographic separation was performed on a µPAC™ (50 cm, PharmaFluidics) column or nanoEase M/Z Peptide BEH C18 Column (130Å, 1.7 µm, 75 µm X 150 mm, Waters, Germany) developing a two-step solvent gradient ranging from 2 to 20% over 47 min and 20 to 32% ACN in 0.08% formic acid within 15 min, at a flow rate of 250 nL/min.

For both, DIA and DDA experiments on the Orbitrap Exploris FAIMS was operated at -50 CV. In DDA LC-MS/MS experiments, full MS data were acquired in the range of 370-1200 m/z at 120.000 resolution. The maximum automatic gain control (AGC) and injection time was set to 3e6 and automatic maximum injection time. Multiply charged precursor ions (2-5) were isolated for higher-energy collisional dissociation (HCD) MS/MS using a 2 Th wide isolation window and were accumulated until they either reached an AGC target value of 2e5 or a maximum injection time of 118 ms. MS/MS data were generated with a normalized collision energy (NCE) of 34, at a resolution of 60.000, with the first mass fixed to 100 m/z. Upon fragmentation precursor ions were dynamically excluded for 120 s after the first fragmentation event.

DIA experiments were acquired in the range of 400–800 m/z at a resolution of 45.000. The AGC was set to 2e5 and the maximum injection time was automatically determined for each scan. DIA windows were acquired with a 5 Th isolation windows (5 Th windows, 1 Th overlap) with stepped NCE 35, 37.5 and 45 at compensation voltages of -50V.

Q Exactive HF-X data was acquired identical as described for the Orbitrap Exploris without the FAIMS Pro interface.

### Data analysis

Reporter ion quantification was performed within the Proteome Discoverer environment (version 2.3.0.484) using the in-house developed, freely available PD node “IMP-Hyperplex” (pd-nodes.org) with a reporter mass tolerance of 10 ppm. The software extracts raw reporter ion intensities from respective spectra for quantification.

Peptide identification was performed using the standard parameters in SpectroMine™ 2.0 against the human reference proteome sequence database (UniProt; version:2018-11-26 accessed April 2019) and yeast reference proteome sequence database (Uniprot; version: 2019-07-25; accessed November 2019). TMT spectral libraries were generated from the DDA files with the highest input and adapted a customized script, kindly provided by Oliver Bernhard from Biognosys. Identification-dependent data aggregation was performed using standard parameters via Spectronaut™ or SpectroMine™ for DIA or DDA, respectively. By default, global median normalization is performed across all experiments. Reporter ion intensities were directly imported into R for further processing. PSMs were filtered to unique peptides using the best scoring (Q-value) PSM for subsequent analysis. Venn Diagrams are based on unique peptide sequences and are calculated using BioVenn.^20^

Identification-independent retention time alignment was based on eight doubly charged precursors (466.7743, 660.3674, 464.779, 437.2732, 587.3539, 587.864, 706.4723) observed in all runs prior to identification-independent data aggregation using the dual indexing approach described above. In detail, in DDA and DIA individual MS/MS scans were indexed according to isolation window and RT. This dual indexing approach for DIA results in a complete grid with no missing data, for DDA we binned precursors within the isolation window (1 Th) and 2 times the FWHM (14 sec) and matched them with +/-1 bin tolerance, to establish a similar grid. Multi-batch TMT identification-independent datasets were normalized by scaling to equal signals per channel. If indicated missing value imputation was performed based on random numbers from normal distribution shifted into the noise by 1.8 in log_10_ space (Lazar et al., 2016). Principal component analysis was internally evaluated considering intra- and inter cluster dimensions using the Davies-Bouldin (DB) index 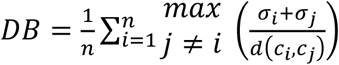. Where n is the number of clusters, c_i_ is the centroid of cluster i, o_i_ is the average Euclidean distance of all elements in cluster i to c_i_. d(c_i_, c_j_) is the distance between centroids of the clusters c_i_ and c_j_.^21^

For both, identification dependent and independent analysis a ComBat based batch correction was performed within the R environment using the sva package.^22,23^

FPKM values were extracted directly from the publicly available ENCODE project (GEO accession: GSE33480) from HeLa and K562 datasets, from which fold changes were calculated. For both DIA and DDA data, loadings of the displayed principal components in Fig 2B were extracted, respective MS/MS scans were intersected with the SpectroMine™/Spectronaut™ derived identifications. PSMs were grouped according to their Master Protein Accession (reporter ion intensity mean) and fold changes between the proteins were calculated. Using the Uniprot database, protein accessions were converted to gene names, which were then intersected with the transcriptome derived FPKM fold changes. Remaining proteins/transcripts were plotted according to their transcript fold change and top 50 (DDA) or top 300 (DIA) protein fold changes are indicated with the respective color. Gene names are indicated for the top 20 (DDA) or top 30 (DIA) transcripts.

## Data Availability

All mass spectrometry-based proteomics data have been deposited to the ProteomeXchange Consortium via the PRIDE partner repository with the dataset identifier [PXD023574].

All R-scripts are deposited via GitHub [DIA-TMT].

## Acknowledgements

We thank all members of our laboratories for helpful discussions. We specifically thank Florian Stanek, Gerhard Dürnberger and Andé Lüttig for bioinformatics support. We thank Oliver Bernhardt and Lynn Verbeke from Biognosis™ for software support. We want to especially thank Rebecca Beveridge, Johannes Griss, Markus Hartl and Elisabeth Roitinger for critical input on the manuscript. This work has been supported by EPIC-XS, project number 823839, funded by the Horizon 2020 program of the European Union and the Austrian Science Fund by ERA-CAPS I 3686-B25-MEIOREC international project.

## Author contributions

J.S. and C.C. conceptualized the project, interpreted data and wrote the manuscript. G.K., K.S. and C.C. optimized equipment and acquired data. C.C. performed sample preparation and data analysis. K.M., J.P., S.M. and J.S. supervised experiments. All authors read and approved the manuscript.

## References

1. Budnik, B., Levy, E., Harmange, G. & Slavov, N. SCoPE-MS: mass spectrometry of single mammalian cells quantifies proteome heterogeneity during cell differentiation. Genome Biol. 19, (2018).

2. Specht, H. et al.. Single-cell proteomic and transcriptomic analysis of macrophage heterogeneity using SCoPE2. Genome Biol. 22, 50 (2021).

3. Izar, B. et al.. A single-cell landscape of high-grade serous ovarian cancer. Nat. Med. 26, 1271–1279 (2020).

4. Miller, A. J. et al.. In Vitro and In Vivo Development of the Human Airway at Single-Cell Resolution. Dev. Cell 53, 117-128.e6 (2020).

5. Zhang, M. J., Ntranos, V. & Tse, D. Determining sequencing depth in a single-cell RNA-seq experiment. Nat. Commun. 11, 774 (2020).

6. Thompson, A. et al.. TMTpro: Design, Synthesis, and Initial Evaluation of a Proline-Based Isobaric 16-Plex Tandem Mass Tag Reagent Set. Anal. Chem. 91, 15941–15950 (2019).

7. Brenes, A., Hukelmann, J., Bensaddek, D. & Lamond, A. I. Multibatch TMT Reveals False Positives, Batch Effects and Missing Values. Mol. Cell. Proteomics 18, 1967–1980 (2019).

8. Karpievitch, Y. V., Dabney, A. R. & Smith, R. D. Normalization and missing value imputation for label-free LC-MS analysis. BMC Bioinformatics 13, S5 (2012).

9. Liu, H., Sadygov, R. G. & Yates, J. R. A Model for Random Sampling and Estimation of Relative Protein Abundance in Shotgun Proteomics. Anal. Chem. 76, 4193–4201 (2004).

10. Tyanova, S., Temu, T. & Cox, J. The MaxQuant computational platform for mass spectrometry-based shotgun proteomics. Nat. Protoc. 11, 2301–2319 (2016).

11. Venable, J. D., Dong, M.-Q., Wohlschlegel, J., Dillin, A. & Yates, J. R. Automated approach for quantitative analysis of complex peptide mixtures from tandem mass spectra. Nat. Methods 1, 39–45 (2004).

12. Bruderer, R. et al.. Extending the Limits of Quantitative Proteome Profiling with Data-Independent Acquisition and Application to Acetaminophen-Treated Three-Dimensional Liver Microtissues. Mol. Cell. Proteomics 14, 1400–1410 (2015).

13. Gillet, L. C. et al.. Targeted Data Extraction of the MS/MS Spectra Generated by Data-independent Acquisition: A New Concept for Consistent and Accurate Proteome Analysis. Mol. Cell. Proteomics 11, (2012).

14. Djebali, S. et al.. Landscape of transcription in human cells. Nature 489, 101–108 (2012).

15. Savitski, M. M. et al.. Measuring and Managing Ratio Compression for Accurate iTRAQ/TMT Quantification. J. Proteome Res. 12, 3586–3598 (2013).

16. Paulo, J. A., O’Connell, J. D. & Gygi, S. P. A Triple Knockout (TKO) Proteomics Standard for Diagnosing Ion Interference in Isobaric Labeling Experiments. J. Am. Soc. Mass Spectrom. 27, 1620–1625 (2016).

17. Karp, N. A. et al.. Addressing accuracy and precision issues in iTRAQ quantitation. Mol. Cell. Proteomics MCP 9, 1885–1897 (2010).

18. Schwacke, J. H., Hill, E. G., Krug, E. L., Comte-Walters, S. & Schey, K. L. iQuantitator: A tool for protein expression inference using iTRAQ. BMC Bioinformatics 10, 342 (2009).

19. Tyanova, S. & Cox, J. Perseus: A Bioinformatics Platform for Integrative Analysis of Proteomics Data in Cancer Research. in Cancer Systems Biology 133–148 (Humana Press, New York, NY, 2018). doi:10.1007/978-1-4939-7493-1_7.

20. Hulsen, T., de Vlieg, J. & Alkema, W. BioVenn – a web application for the comparison and visualization of biological lists using area-proportional Venn diagrams. BMC Genomics 9, 488 (2008).

21. Davies, D. L. & Bouldin, D. W. A Cluster Separation Measure. IEEE Trans. Pattern Anal. Mach. Intell. PAMI-1, 224–227 (1979).

22. Leek, J. T., Johnson, W. E., Parker, H. S., Jaffe, A. E. & Storey, J. D. The sva package for removing batch effects and other unwanted variation in high-throughput experiments. Bioinformatics 28, 882–883 (2012).

23. Johnson, W. E., Li, C. & Rabinovic, A. Adjusting batch effects in microarray expression data using empirical Bayes methods. Biostat. Oxf. Engl. 8, 118–127 (2007).

