## Supplemental Figures for "Comparative proteome signatures of trace samples by multiplexed Data-Independent Acquisition"

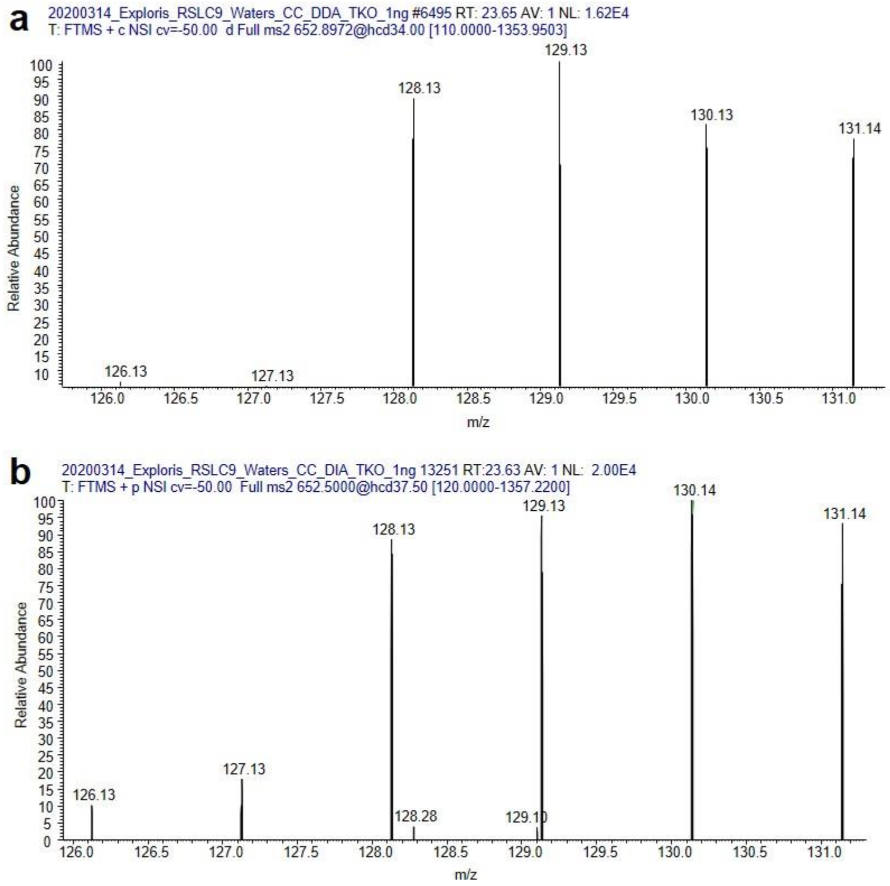


*Supplemental Figure 1: Representative MS/MS scans of met6 knockout TKO11 yeast.* Reporter ion intensities of a representative scan of *met6* knockout yeast at 1 ng total peptide input using both **(a)** DDA and **(b)** DIA acquisition schemes.

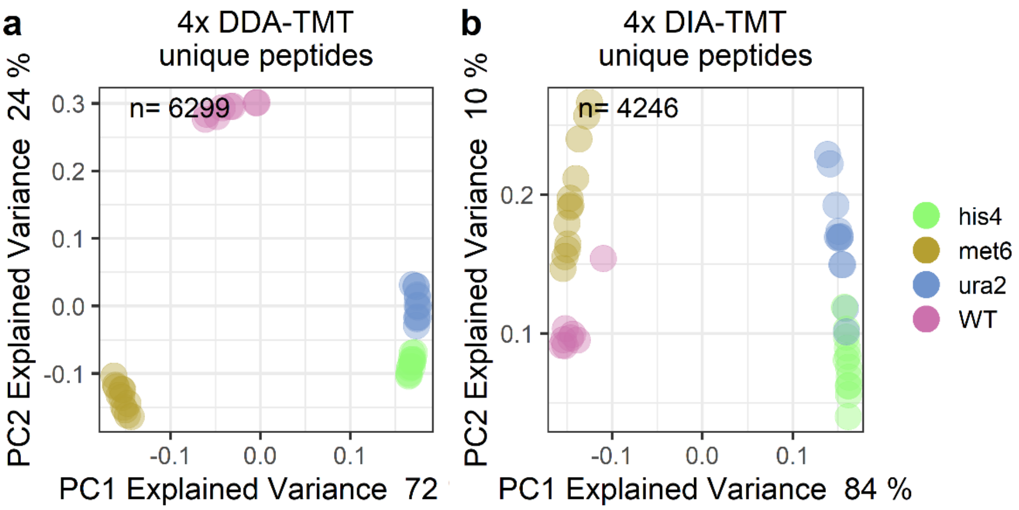


*Supplemental Figure 2: Identification based data aggregation of TKO11 yeast data*. PCA of four analytical runs aggregated via standard identification-dependent method at four peptide inputs (0.5, 1, 5, 10 ng) using **(a)** DDA-TMT or **(b)** DIA-TMT acquisition schemes. Samples are colored according to the respective TKO11 strains (i.e. WT, knockouts: *met6*, *his4*, *ura2*). n= number of unique peptides included in PCA.

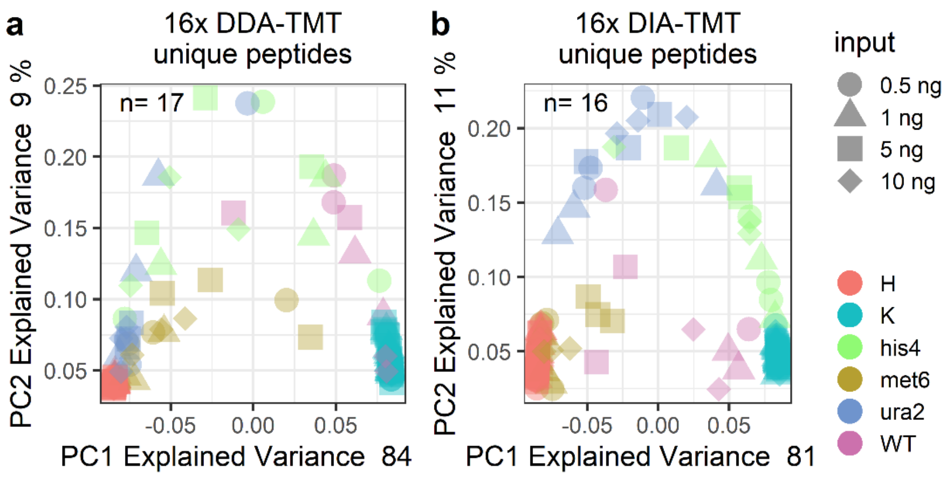


*Supplemental Figure 3: Identification-dependent data aggregation without missing value imputation.* PCA of standard identification-dependent aggregated analytical runs at four peptide inputs (0.5, 1, 5, 10 ng). **(a)** 16 analytical DDA-TMT and **(b)** 16 analytical DIA-TMT runs without missing value imputation. Samples are colored according to channel loadings and the respective peptide input is indicated with different symbols. H = Hela cells, K = K562 cells and the respective TKO11 strains (i.e. WT, knockouts: *met6*, *his4*, *ura2*). n= number of unique peptides included in PCA.
